# The ORF6 accessory protein contributes to SARS-CoV-2 virulence and pathogenicity in the naturally susceptible feline model of infection

**DOI:** 10.1101/2025.04.09.647941

**Authors:** Mohammed Nooruzzaman, Salman L. Butt, Ruchi Rani, Chengjin Ye, Luis Martinez-Sobrido, Ying Fang, Diego G. Diel

**Affiliations:** Department of Population Medicine and Diagnostic Sciences, College of Veterinary Medicine, Cornell University, Ithaca, New York, USA; Texas Biomedical Research Institute, San Antonio, Texas, USA; Department of Pathobiology, College of Veterinary Medicine, University of Illinois at Urbana-Champaign, Urbana, Illinois, USA

## Abstract

Here the infection dynamics, replication, and pathogenicity of a recombinant virus containing a deletion of ORF6 (rWA1ΔORF6) on the backbone of the highly virulent SARS-CoV-2 WA1 virus (rWA1) were investigated and compared to the parental rWA1 virus. While both rWA1 and rWA1ΔORF6 viruses replicated efficiently in cultured cells, the rWA1ΔORF6 virus produced smaller plaques, suggesting reduced cell-to-cell spread. Luciferase reporter assays revealed immune suppressing effects of ORF6 on interferon and nuclear factor kappa B (NF-κB) signaling pathways. Pathogenesis assessment in cats revealed that animals inoculated with rWA1 were lethargic and presented with fever on days 2 and 4 post-infection (pi), whereas rWA1ΔORF6-inoculated animals developed subclinical infection. Additionally, animals inoculated with rWA1ΔORF6 presented reduced infectious virus shedding in nasal and oral secretions and broncho-alveolar lavage fluid when compared with the rWA1-inoculated cats. Similarly, the rWA1ΔORF6-inoculated cats presented reduced virus replication in the respiratory tract as evidenced by lower viral loads and reduced lung inflammation on day 3 and 5 pi when compared to rWA1-inoculated animals. Host gene transcriptomic analysis revealed marked differences in differentially expressed genes (DEG) in the nasal turbinate of animals infected with rWA1 when compared to rWA1ΔORF6. Importantly, type I interferon signaling was significantly upregulated in rWA1ΔORF6 infected cats when compared to rWA1-inoculated animals, which is correlated to the reduced replication of rWA1ΔORF6 in the upper and lower respiratory tracts of infected animals. Collectively, these results demonstrate that the SARS-CoV-2 ORF6 is an important virulence determinant of the virus contributing to the modulation of host antiviral immune responses.

**IMPORTANCE:** SARS-CoV-2 encodes several proteins that inhibit host interferon responses. The accessory protein ORF6 antagonizes interferon signaling by blocking the nucleocytoplasmic trafficking of key transcription factors. In this study, we showed that ORF6 plays an important role in SARS-CoV-2 pathogenesis. While both rWA1 and rWA1ΔORF6 viruses replicated efficiently in cell culture, the rWA1ΔORF6 presented impaired cell-to-spread and reduced innate immune inhibition as compared to the parental rWA1. Pathogenicity study in the feline model revealed an attenuated phenotype of the rWA1ΔORF6 indicating that the ORF6 is a major virulence determinant of SARS-CoV-2. These results also demonstrate that ORF6 contributed to the feline host range of SARS-CoV-2.

## INTRODUCTION

The coronavirus disease 2019 (COVID-19) pandemic, caused by the severe acute respiratory syndrome coronavirus 2 (SARS-CoV-2), emerged in Wuhan, China in late 2019. It has caused more than 777 million human infections with more than 7 million deaths reported to the World Health Organization as of April 8, 2025 (1). Despite significant breakthroughs in vaccine, immunotherapeutic, and antiviral research and development, SARS-CoV-2 remains a major global health burden as variants of the virus continue to emerge. The clinical presentation of COVID-19 varies from asymptomatic infections to severe clinical disease and death, which usually affects individuals with other underlying conditions (e.g., immunosuppression, cardiac diseases, cancers, etc.) and involves imbalanced host innate immune responses (cytokine storm) (2).

Optimal induction of host type I interferon (IFN-I) responses is crucial to COVID-19 clinical outcomes as dysregulated activation of IFN-I signaling pathways can be life-threatening (3, 4). IFN-I responses initiate through recognition of SARS-CoV-2 and its replication products (e.g. ssRNA, dsRNA) through pattern recognition receptors (PRRs) (5). Multiple cytosolic PRRs recognize SARS-CoV-2 RNA (RIG-I, MDA5, TLR3) and host mitochondrial DNA (STING) and activate innate antiviral signaling cascades (6–10). Most of these signaling pathways converge and catalyze the phosphorylation and nuclear translocation of key transcription factors such as interferon regulatory factor 3 (IRF3), IRF7, and nuclear factor kappa B (NF-κB) and the subsequent transcription of IFN-I and IFN-III and proinflammatory cytokines. The secreted IFNs bind to their receptors to activate Janus kinase/signal transducers and activators of transcription (JAK-STAT) signaling pathway to drive the expression of IFN-stimulated genes (ISGs). These ISGs initiate an antiviral state in infected and bystander cells by suppressing critical steps of the viral replication cycle to impair virus spread. Additionally, the expression of ISGs can lead to the activation of immune cells and induce apoptosis of infected cells (11). To counteract host innate antiviral immunity, SARS-CoV-2 has evolved a plethora of mechanisms to inhibit IFN induction and subsequent signaling events. Several viral proteins, such as non-structural protein NSP1, NSP13, NSP14, and accessory ORF3a, ORF6 and ORF9b proteins have been shown to contribute to innate immune evasion (12–22).

SARS-CoV-2 accessory protein ORF6 is uniquely encoded by members of the *Sabrbecovirus* subgenus of Betacoronaviruses. It is a polypeptide with 61 amino acids in length, which functions primarily to antagonize the host interferon signaling pathways, thus limiting host innate immune responses to infection (13–17). The ORF6 protein interacts directly with nucleopore complex components nucleoporin 98 (NUP98) and ribonucleic acid export 1 (RAE1), which suppresses the nuclear import of key transcriptional molecules such as IRF3, STAT1, and STAT2, and inhibit the export of ISG mRNA (13, 16, 17, 20, 23, 24). A recent study showed that ORF6 protein also interacts with RIG-I and inhibits early interferon induction (14).

A hallmark of SARS-CoV-2 is its ability to infect and replicate in a diverse group of animal species. While the function of ORF6 on host immune evasion has been investigated *in vitro*, the studies on the role of ORF6 in SARS-CoV-2 pathogenesis are limited to K18 human angiotensin-converting enzyme 2 (hACE2) transgenic mouse and hamster models (13, 21). In this study, we investigated the role of ORF6 protein using recombinant viruses with ORF6 deletion. The role of ORF6 protein in virus replication and cell-to-cell spread and innate immune evasion were initially characterized *in vitro*, and the infection dynamics, tissue tropism, pathology, and host transcriptional responses were subsequently assessed using the highly susceptible feline model of infection.

## RESULTS

### The ORF6-deletion virus shows efficient replication but reduced plaque size phenotype *in vitro*

To study the contribution of SARS-CoV-2 accessory protein ORF6 in virus infection, replication and pathogenesis, we generated a recombinant virus deficient in ORF6 using reverse genetics technique (25). Using the rWA1 strain as a backbone, a recombinant SARS-CoV-2 lacking the ORF6 gene (rWA1ΔORF6) was generated and rescued (Fig 1A). Complete genome sequencing of the recombinant virus confirmed ORF6 gene deletion and lack of unintended mutations in viral genomes. We next performed multi-step growth kinetics analysis to assess the replication efficiency of both rWA1 and rWA1ΔORF6 viruses using Vero E6 TMPRSS2, Calu-3, and feline lung cells stably expressing cat ACE2 receptor (Feline lung cACE2). Both rWA1 and rWA1ΔORF6 viruses showed comparable replication kinetics in all three cell lines (Fig. 1B), indicating that deletion of ORF6 does not affect virus replication *in vitro*. Next, we assessed the effect of ORF6 deletion on cell-to-cell spread of the virus using plaque assay in Vero E6 cells. The rWA1ΔORF6 virus produced smaller (*p* ≤ 0.0001) plaque size as compared to the parental rWA1 (Fig. 1C and Fig. 1D), suggesting a reduced ability to spread from cell-to-cell of rWA1ΔORF6. These results demonstrate that ORF6 deletion from the SARS-CoV-2 genome does not affect virus replication but leads to reduced cell-to-cell spread *in vitro*.

**FIG 1.**
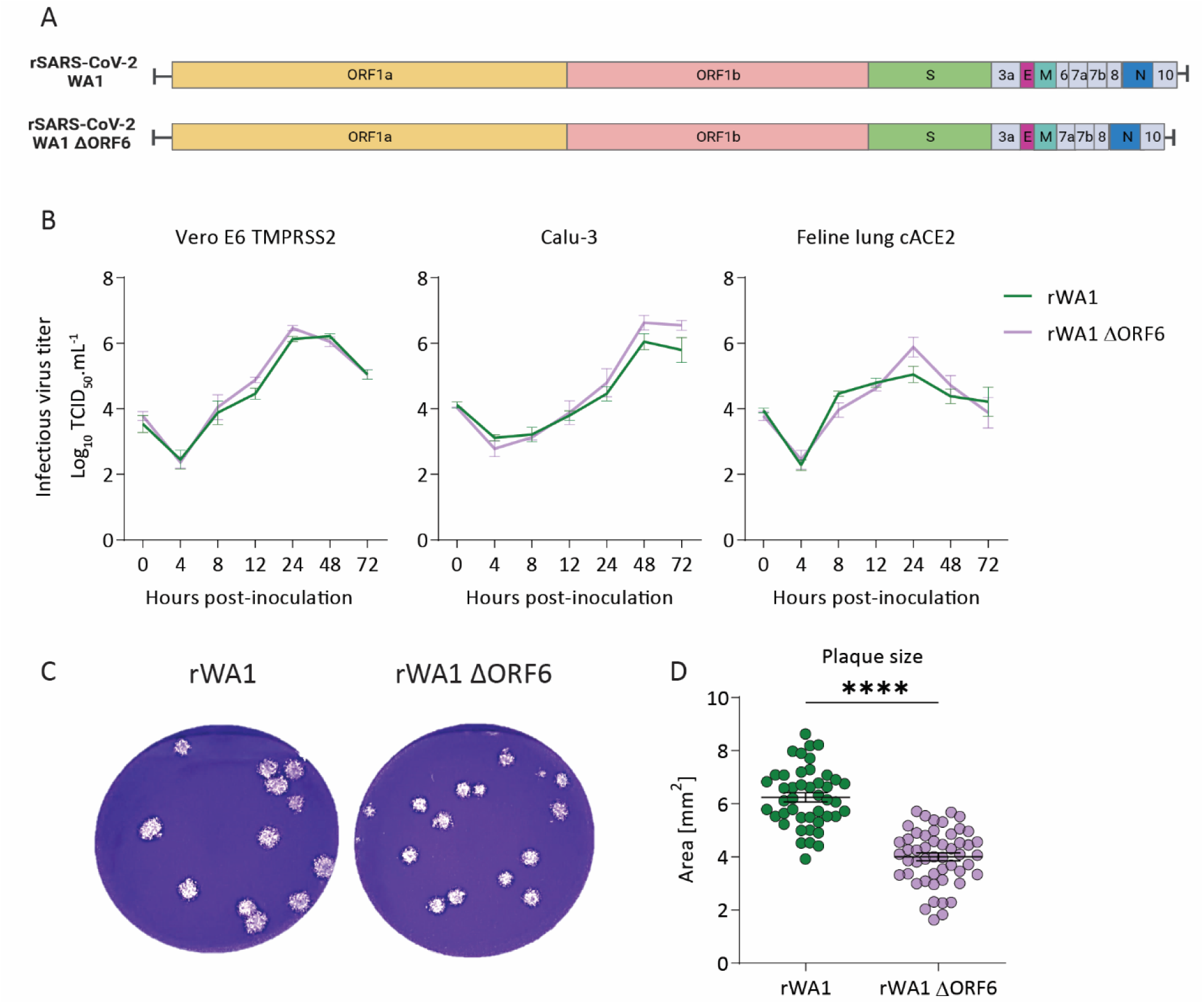
Role of SARS-CoV-2 accessory protein ORF6 in virus replication and spread *in vitro*. (A) Schematic presentation of parental rSARS-CoV-2 WA1 (rWA1) and ORF6 deficient (rWA1ΔORF6) viruses used in this study. (B) Multi-step growth curves. Vero E6 TMPRSS2, Calu-3 and feline lung cACE2 cells were infected (MOI 0.1) with rWA1 and rWA1ΔORF6 viruses and incubated at 4 °C for 1 hour (virus adsorption) and then transferred and incubated at 37°C. Cells were harvested at indicated time points and virus titers were determined by limiting dilutions and expressed as TCID_50._mL^-1^. The limit of detection (LOD) for infectious virus titration was 10^1.05^ TCID_50_.mL^-1^. Data is the mean ± SEM of three independent experiments. (C) Plaque phenotype. Vero E6 cells were infected with rWA1 and rWA1ΔORF6 and overlaid with media containing 0.5% agarose. Plates were incubated at 37 °C for 72 h, the agarose overlay was removed, cells were fixed, and the monolayer was stained with 0.5% crystal violet. (D) Viral plaque sizes. The diameters of viral plaques were measured using a scale in millimeters. Data indicate means ± SEM. Mann Whitney U test, **** *p* ≤ 0.0001.

### SARS-CoV-2 accessory protein ORF6 inhibits multiple innate immune pathways *in vitro*

We investigated the role of SARS-CoV-2 ORF6 protein in modulating the host innate antiviral pathways *in vitro* by using luciferase reporter assays. For this, we evaluated the inhibitory effect of ORF6 on activation of IFN-β-, IRF3- and NF-κB-driven promoters. Additionally, we investigated the inhibitory effect of ORF6 on two positive regulatory domains (PRDs) of the IFN-β enhanceosome, PRDII and PRDIII, that are recognized by transcription factors NF-κB and IRF3/IRF7, respectively (26). For this, we co-transfected HEK293T cells with plasmids encoding ORF6 protein or an empty vector, with IFN-β-, IRF3-, NF-κB-, PRDII-, and PRDIII-driven Firefly luciferase reporter plasmids. A plasmid constitutively expressing Renilla luciferase reporter gene (pRN-Luc) was used as a control to normalize transfection efficiencies. At 24 hours post-transfection, cells were stimulated with Sendai virus (SeV) or poly(I:C) (two interferon inducers), or TNF-α (a NF-κB pathway inducer). After 12 hours of stimulation, Firefly luciferase activity was quantified to assess the inhibitory role of ORF6 protein on these innate signaling pathways. Overexpression of ORF6 protein significantly downregulated activation of poly(I:C)- and SeV-induced IFN-β signaling (Fig. 2A), SeV-induced IRF3 signaling (Fig. 2B), TNF-α-induced NF-κB signaling (Fig. 2C), SeV- and TNF-α-induced PRDII signaling (Fig. 2D), and poly(I:C)- and SeV-induced PRDIII signaling pathways (Fig. 2E). These results demonstrate that ORF6 inhibits both the IFN-β and NF-κB signaling pathways.

**FIG 2.**
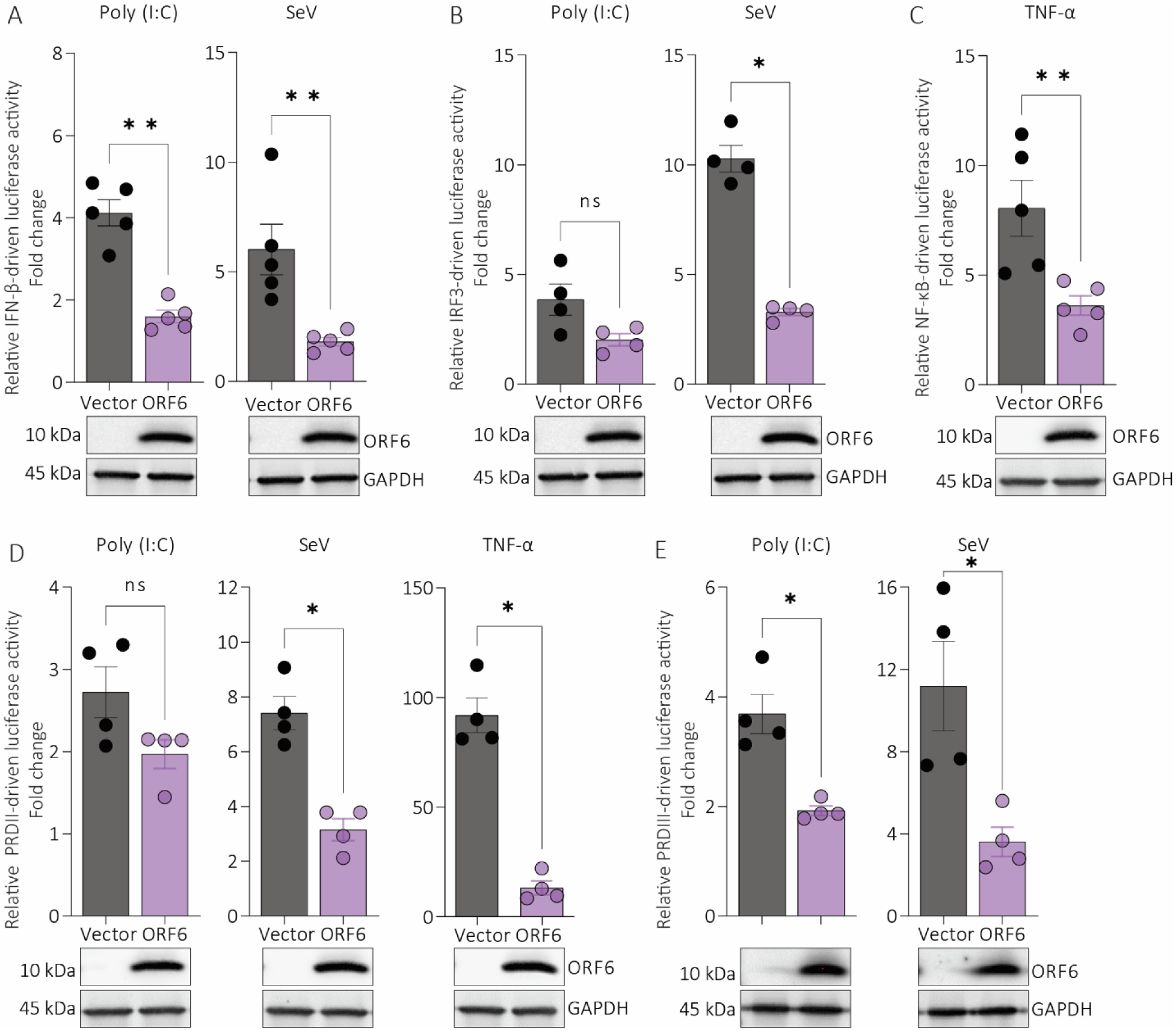
Innate immune inhibition by SARS-CoV-2 accessory protein ORF6. HEK293T cells were transfected with plasmids encoding ORF6 or an empty vector and with IFN-β (A), IRF3 (B), NF-κB (C), PRDII (D), and PRDIII (E) Firefly reporter plasmids. At 24 hours post-transfection, cells were stimulated with poly(I:C), SeV or TNF-α for 12 hours. Cell lysates were harvested, and Firefly luciferase activity was measured using a luminometer. The ratio of luminescence obtained from target reporters (IFN-β-Luc, IRF3-Luc, NF-κB-Luc, PRDII-Luc, and PRDIII-Luc) to luminescence from the control Renilla reporter (pRN-Luc) was calculated to normalize for transfection efficiencies. The relative IFN-β-, IRF3-, NF-κB-, PRDII-, or PRDIII-driven luciferase activity was calculated as fold change over the unstimulated cells. Data indicates mean ± SEM of 3-4 independent experiments. Mann-Whitney U test, * *p* ≤ 0.05, ns = not significant.

Next, we tested the relevance of the innate immune inhibitory effects of ORF6 overexpression in the context of SARS-CoV-2 infection. For this, HEK293T cells expressing the human ACE2 receptor (HEK293T-hACE2) were transfected with IFN-β- or NF-κB-firefly luciferase plasmids for 6 hours. Cells were then infected with rWA1 or rWA1ΔORF6 (MOI 3) for 12 hours followed by stimulation with SeV or TNF-α for 8 hours. Cell lysates were harvested, and the luciferase activity was quantified as above. Infection of HEK293T-hACE2 cells with parental rWA1 or rWA1ΔORF6 viruses induced low to moderate IFN-β or NF-κB reporter activation (Fig. 3A-Fig.3C). However, subsequent stimulation of the SARS-CoV-2 infected cells with SeV and TNF-α greatly enhanced the IFN-β and/or NF-κB promoter signaling which was comparable or higher than the SeV or TNF-α stimulation alone (Fig. 3A-Fig.3C). Of note, SeV stimulation of rWA1ΔORF6 infected cells induced higher IFN-β promoter activation (p<0.05) than the rWA1 infected cells (Fig. 3A), suggesting that deletion of ORF6 from rWA1 virus resulted in impaired ability of the virus to inhibit interferon signaling pathway. Similarly, TNF-α stimulation of the rWA1ΔORF6 infected cells also resulted in higher NF-κB promoter activation (p<0.05) as compared to TNF-α stimulated rWA1 infected cells (Fig. 3C). Collectively, our luciferase reporter assays using both plasmid overexpression and recombinant viruses highlight the important role of ORF6 on innate immune inhibition during SARS-CoV-2 infection.

**FIG 3.**
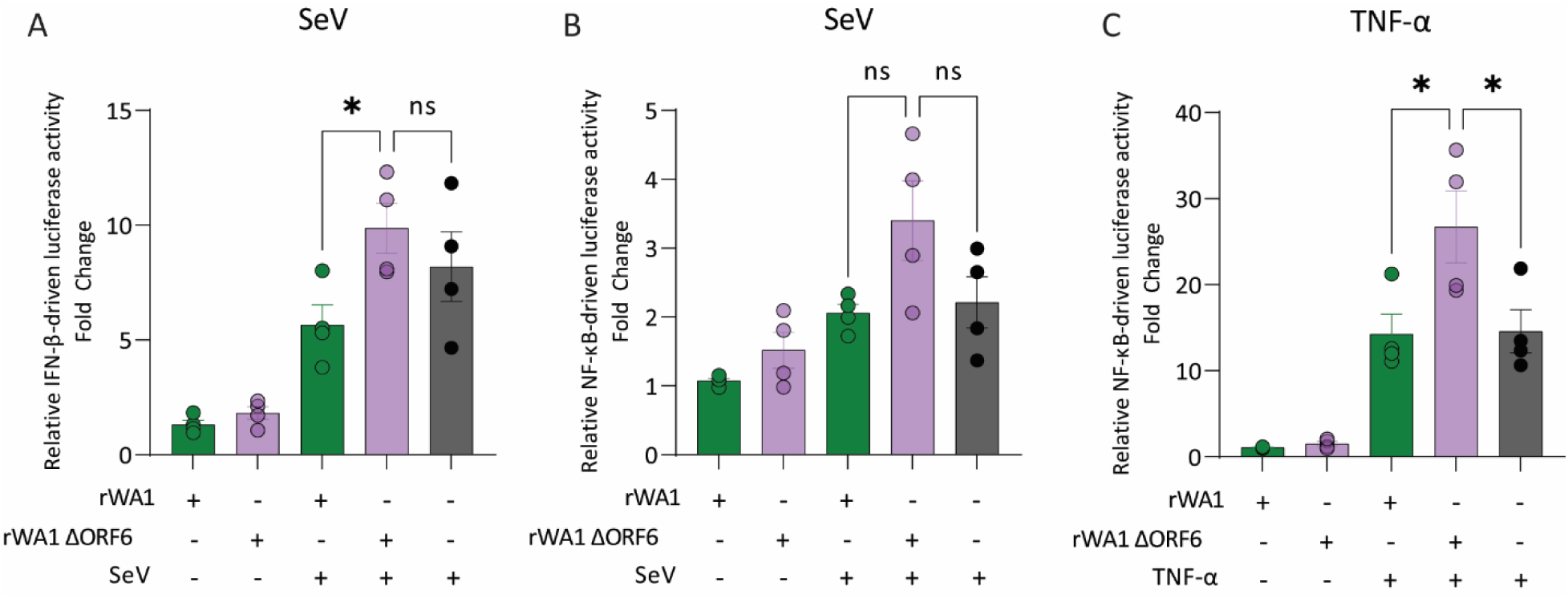
Role of ORF6 on innate immune inhibition during SARS-CoV-2 infection *in vitro*. (A) HEK293T-hACE2 cells were transfected with IFN-β (A) or NF-κB (B-C) Firefly reporter plasmids for 6 hours. Cells were then infected with rWA1 or rWA1ΔORF6 (MOI = 3) for 12 hours. After that, cells were stimulated with SeV or TNF-α for 8 hours, cell lysates were harvested, and the luciferase activity was measured using a luminometer. The ratio of luminescence obtained from the target reporters (IFN-β-Luc or NF-κB-Luc) to luminescence obtained from the control Renilla reporter (pRN-Luc) was calculated to normalize the transfection efficiency. The relative IFN-β or NF-κB promoter driven luciferase activity was expressed as fold change over the unstimulated cells. Data indicates mean ± SEM of 4 independent experiments. One-way ANOVA with multiple comparison test, * *p* ≤ 0.05, ns = not significant.

### Infection with rWA1ΔORF6 leads to subclinical infection and reduced virus shedding in cats

Three groups of experimental cats were inoculated with rWA1 or rWA1ΔORF6 viruses, or mock-infected (Fig. 4A). Clinical signs, virus replication in tissues, virus shedding and pathology in respiratory tissues were evaluated (Fig. 4A). Cats inoculated with rWA1 were depressed and presented elevated body temperature throughout the 5-day experimental period with significant rise in the body temperature being observed on day 2 post-infection (pi) (*p* ≤ 0.05) when compared to rWA1ΔORF6- and mock-inoculated groups (Fig. 4B). No clinical sign was observed in rWA1ΔORF6- and mock inoculated cats. Both virus-inoculated groups lost or maintained body weight, while control cats gained body weight throughout the study period (Fig. 4C).

**FIG 4.**
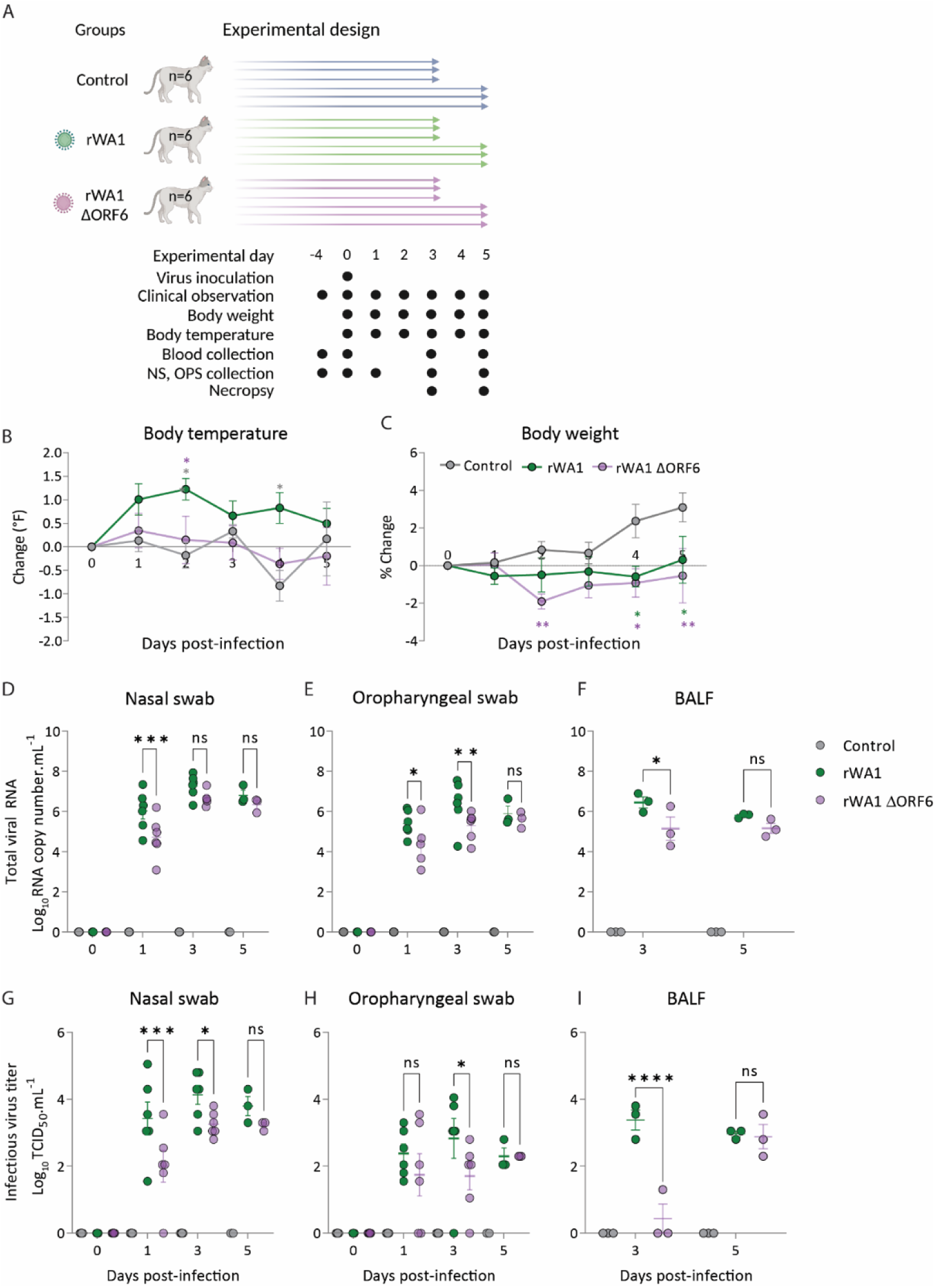
ORF6 contributes to SARS-CoV-2 virulence and pathogenesis in cats. (A) Experimental study design. (B) Changes (°F) in body temperature and (C) body weight (%) in cats intranasally inoculated with rWA1 and rWA1ΔORF6. SARS-CoV-2 RNA loads quantified by rRT-PCR in nasal (D) and oral (E) secretions collected on days 0, 1, 3 and 5 pi and in bronchoalveolar lavage fluid (F) collected on days 3 and 5 pi. Infectious SARS-CoV-2 loads in nasal (G) and oral (H) secretions and BALF (I) were determined by virus titration in rRT-PCR-positive samples. Virus titers were determined using endpoint dilutions and expressed as TCID_50_.mL^-1^. The limit of detection (LOD) for infectious virus titration was 10^1.05^ TCID_50_.mL^-1^. Data indicates mean ± standard deviation of 3-6 animals per group per time point. 2-way ANOVA with multiple comparison test, * *p ≤* 0.05; ** *p ≤* 0.01; *** *p ≤* 0.005; **** *p ≤* 0.0001.

We next assessed the dynamics of virus shedding in inoculated cats. To this end, nasal (NS) and oropharyngeal (OPS) swabs were collected from control and inoculated cats on days 0, 1, 3 and 5 pi and bronchoalveolar lavage fluid (BALF) was collected on days 3 and 5 pi at necropsy and tested by rRT-PCR and endpoint virus titrations. Overall, viral RNA loads in NS, OPS and BALF were lower in rWA1ΔORF6 inoculated cats when compared to rWA1 inoculated animals throughout the experiment. The rWA1ΔORF6 inoculated cats presented significantly lower viral RNA in nasal swabs on day 1 pi (*p* ≤ 0.001) than rWA1 inoculated animals (Fig. 4D). Similarly, lower viral RNA levels were found in OPS of rWA1ΔORF6 inoculated cats when compared to rWA1 inoculated animals on days 1 (*p* ≤ 0.05) and 3 (*p* ≤ 0.01) pi (Fig. 4E). Consistent with virus replication in the upper respiratory tract, lower viral RNA loads were detected in BALF of rWA1ΔORF6 inoculated cats when compared to rWA1 inoculated animals on day 3 pi (*p* ≤ 0.05) (Fig. 4F).

To study the dynamics of infectious virus shedding in NS, OPS and BALF, we quantified infectious viruses in rRT-PCR-positive swab samples by endpoint virus titrations. Similar to the rRT-PCR results, infectious virus titers in NS, OPS and BALF were lower in rWA1ΔORF6 inoculated cats than in rWA1 inoculated animals (Fig. 4G-4I). Infectious virus titers in NS were significantly lower in rWA1ΔORF6 inoculated cats on days 1 (0-2.55 Log TCID_50_.mL^-1^ vs 1.55-5.05 Log TCID_50_.mL^-1^, *p* ≤ 0.001) and 3 pi (2.8-3.8 Log TCID_50_.mL^-1^ vs 3.05-4.8 Log TCID_50_.mL^-1^, *p* ≤ 0.05) when compared to rWA1 inoculated animals (Fig. 4G). The infectious virus titers in OPS were about 1 Log lower than virus titers obtained in NS. Similarly, rWA1ΔORF6 inoculated cats had a lower viral load in OPS on day 3 pi (*p* ≤ 0.05) than rWA1 inoculated animals (Fig. 4H). In BALF, infectious virus was isolated from all three cats inoculated with rWA1 virus on day 3 pi, while only one cat from rWA1ΔORF6 inoculated group presented infectious virus in BALF (Fig. 4I). The BALF from all inoculated cats on day 5 pi showed positive virus isolation and a comparable virus titer between rWA1 and rWA1ΔORF6 inoculated animals (Fig. 4I). Together, these results demonstrate that deletion of ORF6 from SARS-CoV-2 WA1 genome resulted in subclinical infection and reduced virus replication and shedding in cats. Thus, ORF6 contributed to SARS-CoV-2 virulence and pathogenesis in the naturally susceptible feline model of SARS-CoV-2 infection.

### Deletion of ORF6 from SARS-CoV-2 WA1 results in limited virus replication in respiratory tissues

We studied the effect of ORF6 deletion on tissue tropism and replication of SARS-CoV-2. For this, tissues collected from rWA1- and rWA1ΔORF6-inoculated cats were tested for the virus presence by titrations and *in situ* hybridization (ISH). Tissue samples, including nasal turbinate, tonsil/palate, retropharyngeal lymph node, trachea, lungs, mediastinal lymph node, heart, liver, spleen, kidney, intestine, and mesenteric lymph node, were collected at necropsy on days 3 and 5 pi from control and inoculated cats and processed for infectious virus quantification. The highest infectious virus titers were detected in nasal turbinates from rWA1 inoculated cats on day 3 pi (titers ranging 5.55-6.05 Log_10_ TCID_50_.mL^-1^), while significantly lower virus titers were obtained from rWA1ΔORF6 inoculated cats (titers ranging 1.8-5.05 Log^10^ TCID_50_.mL^-1^) (*p* ≤ 0.05) (Fig. 5A). Similarly, on day 5 pi, rWA1ΔORF6 inoculated cats had lower virus titers in nasal turbinates (titer ranging 4.3-5.8 Log_10_ TCID_50_.mL^-1^) than rWA1 inoculated cats (titers ranging 3.3-4.8 Log_10_ TCID_50_.mL^-1^) (Fig. 5B). The rWA1ΔORF6 inoculated cats also had significantly lower virus titers in trachea (*p* ≤ 0.05) and lungs (*p* ≤ 0.05) on day 3 pi and tonsil/palate (*p* ≤ 0.05) and trachea (*p* ≤ 0.001) on day 5 pi when compared to rWA1 inoculated cats (Fig. 5A and Fig. 5B). No infectious virus was detected in tissues collected from the control cats.

**FIG 5.**
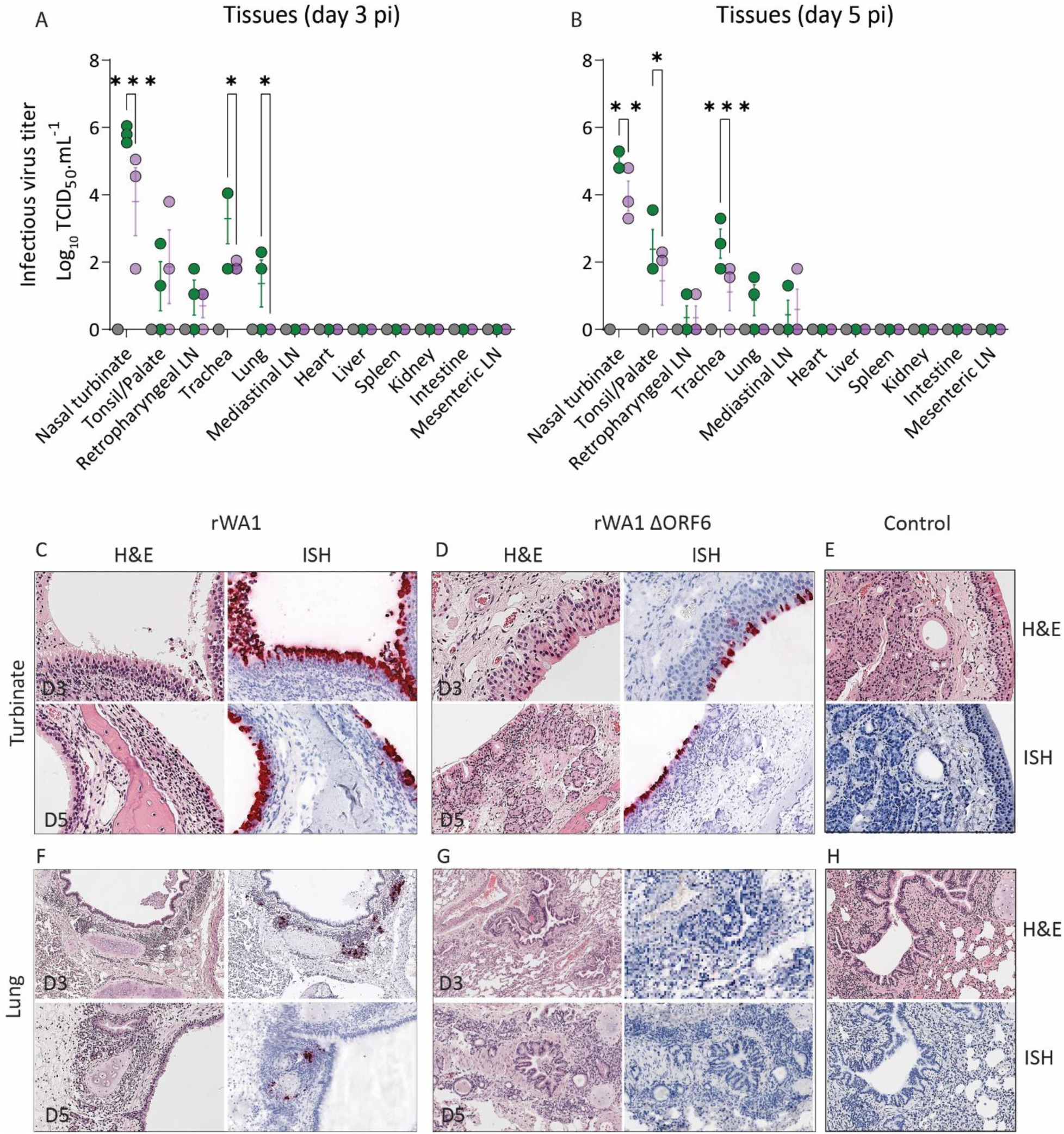
rWA1ΔORF6 presents reduced replication and pathology in the respiratory tract of cats. Infectious SARS-CoV-2 in tissues assessed by virus titration in rRT-PCR positive tissue samples collected on days 3 (A) and 5 (B) pi. Virus titers were determined using endpoint dilutions and expressed as TCID_50_.mL^-1^. The limit of detection (LOD) for infectious virus titration was 10^1.05^ TCID_50_.mL^-1^. Data indicates mean ± standard deviation of 3 animals per group per time point. 2-way ANOVA with multiple comparison test, * *p ≤* 0.05; *** *p ≤* 0.005. Hematoxylin and eosin (H&E) staining and *in situ* hybridization (ISH) in nasal turbinate (C-E) and lung (F-H) of cats inoculated with rWA1 and rWA1ΔORF6 and control cats. Nasal turbinate and lungs were collected from rWA1 (C and F) and rWA1ΔORF6 (D and G) or mock (E and H) inoculated cats on days 3 and 5 pi. In ISH, the viral RNA (red labeling) was performed using a probe targeting the SARS-CoV-2 S RNA.

We next localized virus replication sites in tissues of inoculated cats using *in situ* hybridization (ISH). For this, we stained nasal turbinate, trachea and lungs collected from all animals on days 3 and 5 pi for SARS-CoV-2 RNA using RNAscope^®^ ZZ technology. The epithelial cells of nasal mucosa were the predominant cell types showing positive labeling for SARS-CoV-2 RNA with intense hybridization being observed on day 3 pi, which was slightly less abundant on day 5 pi. The nasal turbinates collected from rWA1 inoculated cats showed more intense labeling for SARS-CoV-2 RNA on days 3 and 5 pi when compared to rWA1ΔORF6 inoculated animals (Fig. 5C-5D). All SARS-CoV-2 inoculated cats from both groups showed positive viral RNA hybridization in nasal turbinates, however, the intensity and number of positive cells were higher in rWA1 inoculated cats when compared with rWA1ΔORF6 inoculated animals. In the trachea localized viral staining was detected in cells within submucosal interstitial stroma, as well as cells associated with submucosal glandular or vascular elements. Viral RNA hybridization in the trachea was also higher in rWA1 inoculated cats on both days 3 and 5 pi as compared to rWA1ΔORF6 inoculated animals (data not shown). Of note, all 6 cats inoculated with rWA1ΔORF6 showed positive RNA labeling in trachea, while one rWA1 inoculated cat on day 3 was negative. In lungs, intense viral RNA hybridization was mostly observed in bronchial glandular epithelial cells with sparse staining of bronchial epithelial cells of rWA1 inoculated cats on both days 3 (2 cats) and 5 (1 cat) pi (Fig. 5F). No viral RNA labeling was found in lungs of rWA1ΔORF6 inoculated cats on either day 3 or 5 pi (Fig. 5G). Tissues collected from control cats remained negative for viral RNA using ISH (Fig. 5E and Fig. 5H).

### Deletion of ORF6 from SARS-CoV-2 WA1 virus genome results in limited lung pathology in cats

The histological changes in the nasal turbinate, trachea, and lungs of cats were evaluated on days 3 and 5 pi. In nasal turbinate, loss of mucosal epithelial cells and mild to moderate epithelial necrosis were noticed in both rWA1 and rWA1ΔORF6 inoculated cats on days 3 and 5 pi (Fig. 5C-5D). Aggregates of fibrin, necrotic debris, and inflammatory cells in nasal passages were also observed in both inoculated groups. No remarkable histological changes were observed in the trachea. In lungs, bronchiolar necrosis, and mixed inflammation with exudates in the lumen were observed in two rWA1 and rWA1ΔORF6 inoculated cats on day 3 pi and two rWA1 and one rWA1ΔORF6 inoculated cats on day 5 pi (Fig. 5F-5G). Thickening of the alveolar septa due to mononuclear infiltration was noticed in all three rWA1-inoculated cats but in one rWA1ΔORF6-inoculated animal on day 3 pi. Alveolar septal thickening was also noticed in all rWA1 and rWA1ΔORF6 inoculated cats on day 5 pi (Fig. 5F-5G). Exudates in the alveolar lumen were found in most of the inoculated cats on both days 3 and 5 pi, with lower severity in rWA1ΔORF6 inoculated cats. The control cats had normal histological architectures in all tissues studied (Fig. 5E-5H). Collectively, these results demonstrate that the rWA1ΔORF6 virus induces limited pathology in lungs of cats.

### Deletion of ORF6 from SARS-CoV-2 WA1 virus genome results in differential transcriptional profile in nasal turbinate of cats

We performed bulk RNA-seq to investigate the differential expression of genes in nasal turbinates, the first site of SARS-CoV-2 replication and where we observed the highest viral loads, collected from control or from rWA1 and rWA1ΔORF6 infected cats. Differential gene expression analysis revealed several thousand differentially expressed host genes (DEGs). The relative gene expression obtained by comparing rWA1ΔORF6 vs control and rWA1 vs control is summarized in the heatmap showing the top 5000 hierarchically clustered DEGs across the three groups (Fig. 6A). Each column represents the gene expression profile from individual samples and the colored bars (blue, white and red bars represent z score values in the color key) represents up-regulated (red), down-regulated (blue), or unchanged (white) gene expression. A comparison of significant DEGs revealed a total 394 up- and 104 down-regulated genes in rWA1 vs control group (Fig. 6B), 134 up- and 8 down-regulated DEGs in rWA1ΔORF6 vs control (Fig. 6C), and 19 up- and 17 down-regulated DEGs in rWA1 vs rWA1ΔORF6 groups (Fig. 6D).

**FIG 6.**
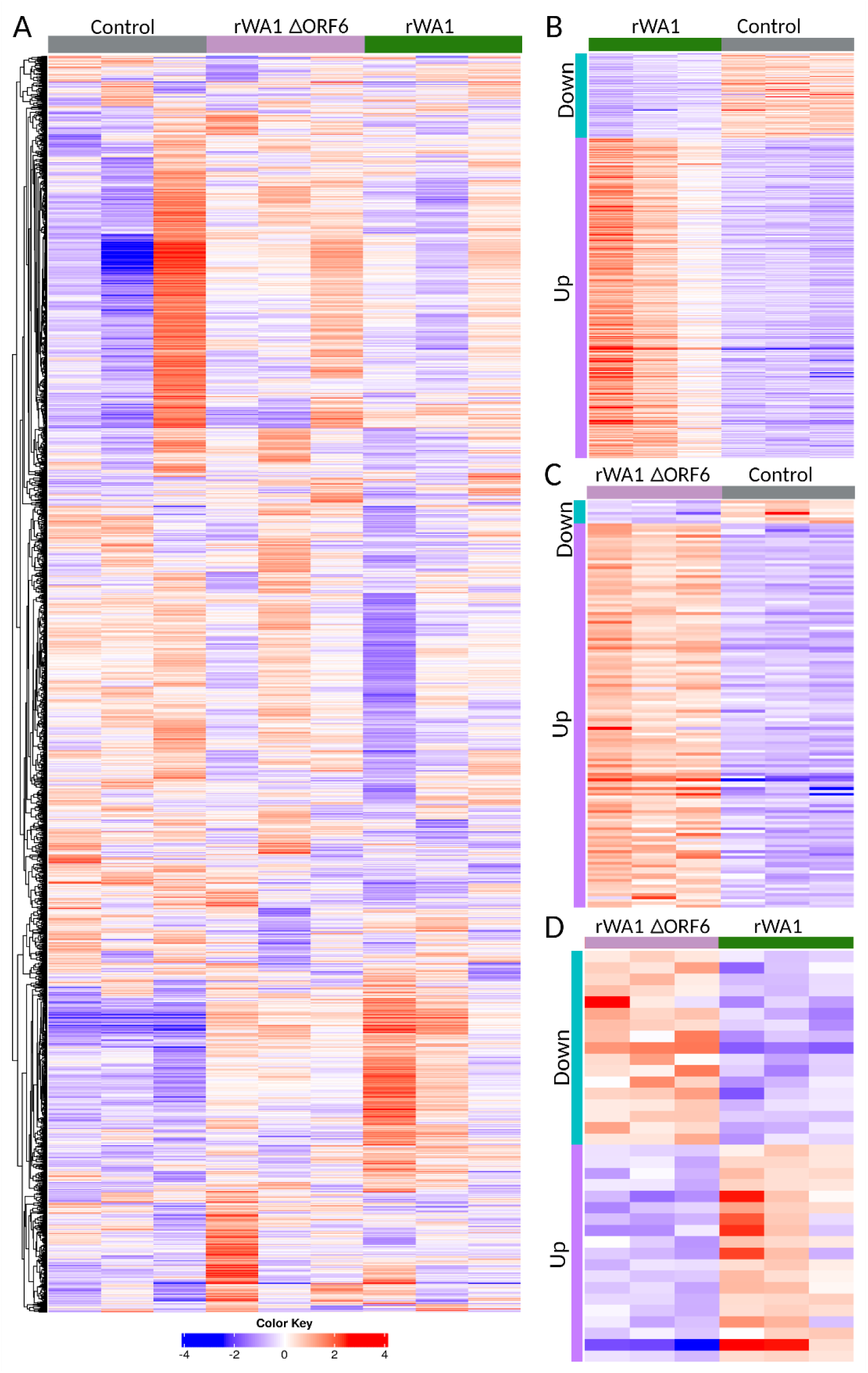
Heatmaps of differential gene expression in nasal turbinate of cats infected with SARS-CoV-2. Nasal turbinate was collected from rWA1 and rWA1ΔORF6 or mock-inoculated cats on day 3 pi and subjected to RNA-Seq analysis. (A) Heatmap of hierarchically clustered differentially expressed genes (DEGs) in nasal turbinate tissues of rWA1, rWA1ΔORF6, and mock-infected (control) cats. Each column represents the gene expression profiles from individual samples across three groups. A total of 5000 genes represented by colored bars represents z-score values in the color key where red represents up-regulation, blue represents down-regulation, and white represents unchanged gene expression. A group comparison of DEGs in rWA1 vs control showed 394 genes up- and 104 down-regulated genes (B) 134 up- and 8 down-regulated genes in rWA1ΔORF6 vs control (C), and 19 up- and 17 down-regulated genes in rWA1 vs rWA1ΔORF6 groups (D). The up- and down-regulated DEGs are clustered across three samples from each group.

### Gene set enrichment analysis (GSEA)

#### Cellular pathway enrichment

The significant DEGs from the three groups (control, rWA1, and rWA1ΔORF6) were further used for pathway enrichment analysis, and the top significantly enriched GO biological pathways are presented as dot plots in Fig. 7A-7C. This analysis revealed that the rWA1 and rWA1ΔORF6 triggered different pathways in the nasal turbinate of infected cats. The significantly enriched pathways were related to innate immune response in both rWA1 and rWA1ΔORF6 infected cats. Overall, the fold enrichment of innate immune response pathways was much higher in rWA1ΔORF6 relative to rWA1 infected cats. In rWA1 infected cats, the antiviral signaling pathways was 12-fold enriched and the stress response pathway was enriched three (03) fold (Fig. 7A). In contrast, in rWA1ΔORF6 infected cats, the type I interferon signaling pathway was enriched 40-fold and the response to type I interferon pathway was enriched 27-fold when compared to rWA1 inoculated cats (Fig. 7B).

**FIG 7.**
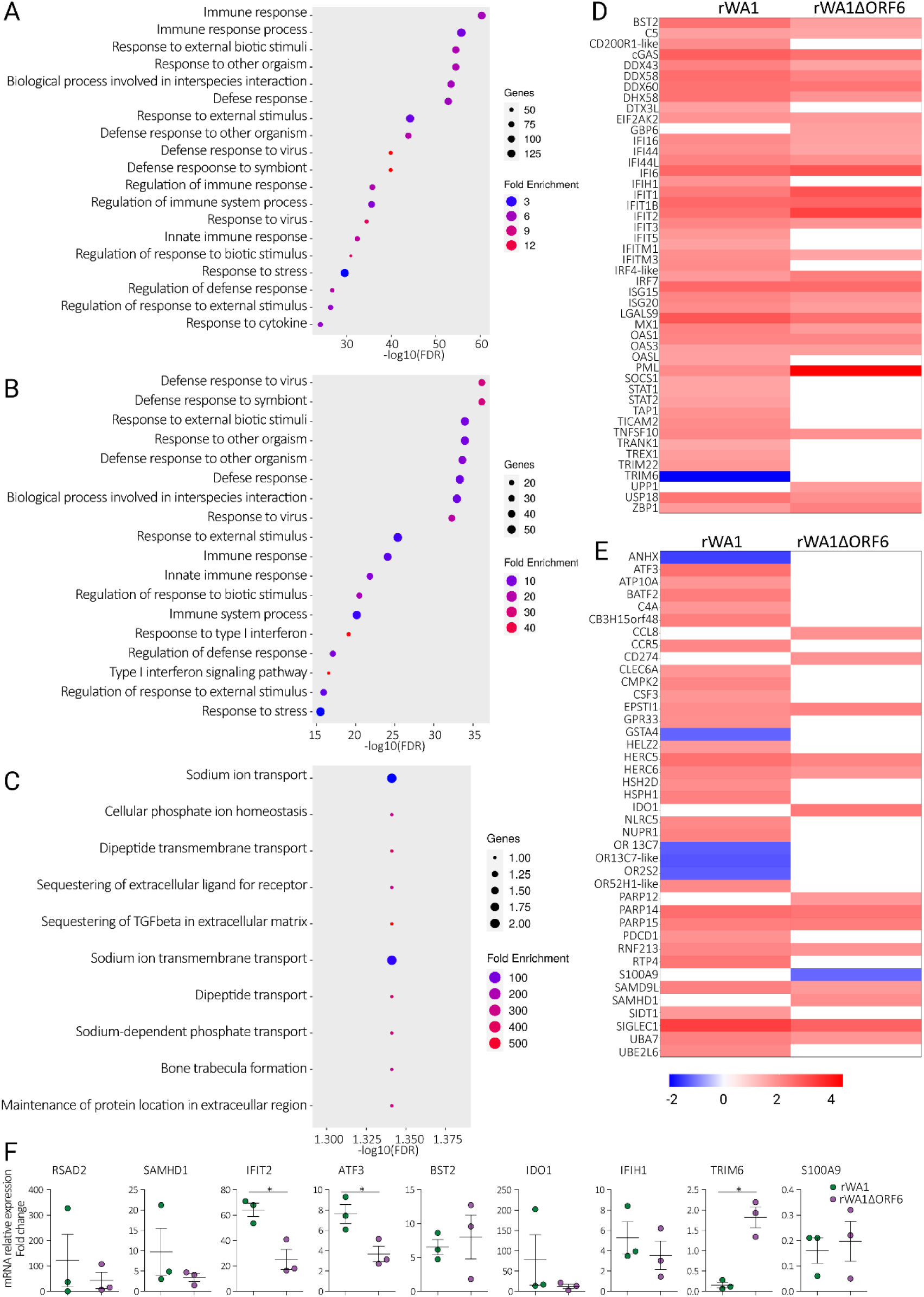
rWA1 and rWA1ΔORF6 trigger different pathways in the nasal turbinate of cats. Nasal turbinate was collected from rWA1 and rWA1ΔORF6 or mock-inoculated cats on days 3 pi and subjected to RNA-Seq analysis. Dot plot showing the upregulated GO biological processes across three group comparisons, rWA1 vs control (A), rWA1ΔORF6 vs control (B), and rWA1ΔORF6 vs rWA1 (C). Heatmaps showing log2FC expression profile of top 40 interferon-stimulated genes (ISGs) (D) and top 40 transcription factors (E) in nasal turbinate of cats infected with rWA1 and rWA1ΔORF6. (F) Dot plots showing verification of gene expression of top 9 differentially expressed genes in nasal turbinate of cats. Data indicate mean ± SEM of 3 animals per group. Mann-Whitney U test, *p* ≤ 0.05 indicates statistical significance.

Next, we investigated the association of the presence or absence of ORF6 in rWA1 and rWA1ΔORF6 infected cats on the host innate immune response, especially on ISGs expression, and their transcription factors. To investigate the differences in innate immune response to rWA1 and rWA1ΔORF6, we selected 80 significant DEGs associated with innate immune response from groups of rWA1 vs control and rWA1ΔORF6 vs control group. Their relative fold changes are summarized in the heatmap presented in Fig. 7D-7E. Notably, ISGs such as BST2, Mx1, IFI6, and cGAS were up-regulated and TRIM6 was downregulated in rWA1 when compared to rWA1ΔORF6 infected cats. Expression of GBP6 and UPP1 was unchanged in rWA1 infected cats, while these genes were up regulated in rWA1ΔORF6 inoculated animals. Similarly, RSAD2, IFIT1, IFIT2, and IRF7 were upregulated, while the expression of several ISGs (IFIH1, IFIT5, IFITM1, IRF4-like, PML, SOCS1, STAT1, STAT2, TRIM22, TRIM6, TRAM1, and TREX1) remained unchanged in rWA1ΔORF6-infected cats, compared to rWA1-infected cats where these genes were upregulated. Among the transcription factors, genes associated with olfactory transduction (OR13C-7, OR2S2), ANHX, and GSTA4 were downregulated in rWA1-infected cats, while their expression remained unchanged in rWA1ΔORF6-infected cats. Additionally, CCL8, CD274, IDO1, PARP12, S100A9, and SAMHD1 expression remained unchanged in rWA1-infected cats, whereas these genes were upregulated in rWA1ΔORF6-infected cats (Fig. 7E).

We selected the top 9 differentially expressed genes from all three comparison panels, rWA1 vs control, rWA1ΔORF6 vs control and rWA1ΔORF6 vs rWA1 for verification of gene expression by rRT-PCR (Fig. 7F). The IFIT2 and ATF3 genes were significantly (*p* ≤ 0.05) upregulated in rWA1 compared to the rWA1ΔORF6 inoculated animals, whereas TRIM6 was significantly (*p* ≤ 0.05) downregulated in rWA1 compared to rWA1ΔORF6 inoculated animals. Other genes such as RSAD2, SAMHD1, BST2, IDO1, IFIH1, and S100A9 were upregulated but remained comparable between both rWA1 and rWA1ΔORF6 inoculated animals (Fig. 7F).

## DISCUSSION

Our study builds on previous work (21), demonstrating that the SARS-CoV-2 accessory protein ORF6 is a bona fide virulence determinant that contributes to disease pathogenesis by modulating host innate immune responses. Using a highly relevant and naturally susceptible feline model of SARS-CoV-2 infection (27–34), we demonstrated that deletion of ORF6 from the SARS-CoV-2 WA1 genome results in an attenuated disease phenotype. The rWA1ΔORF6-infected animals present subclinical infection and markedly reduced virus replication and associated pathology in the upper and lower respiratory tracts.

The SARS-CoV-2 ORF6 protein functions to counteract host innate immune responses by selectively blocking the nuclear import and translocation of IRF3 and STAT transcription factors (13, 16, 17, 23, 24, 35). To further investigate the inhibition of downstream interferon signaling pathways by ORF6 protein, we used luciferase reporter assays. In line with previous studies, transient overexpression of ORF6 protein significantly downregulated Sendai virus (SeV)-induced IFN-β and IRF3 promoter activation (13, 14). However, we also found a significant inhibition of TNF-α induced NF-κB promoter activation upon ORF6 overexpression, which differs from an earlier study (13). We further investigated the IFN-β promoter activity and included two positive regulatory domains (PRDs) of the IFN-β enhanceosome, PRDII and PRDIII, that are recognized by transcription factors NF-κB and IRF3/IRF7, respectively (26). The ORF6 protein also inhibited SeV and TNF-α induced PRDII and poly(I:C) and SeV-induced PRDIII promoter activation. To address the relevance of these findings in the context of infection, we infected HEK293T-hACE2 cells with rWA1 and rWA1ΔORF6 viruses and quantified the activation of both IFN-β and NF-κB reporters. Infection of HEK293T-hACE2 cells with recombinant viruses alone induced low IFN-β and NF-κB activation, which increased significantly upon subsequent stimulation with SeV and/or TNF-α. Of note, compared to the parental rWA1, stimulation of rWA1ΔORF6-infected cells with SeV and TNF-α significantly enhanced IFN-β and NF-κB-promoter activation, respectively. Collectively, these results demonstrate the innate immune inhibition ability of the SARS-CoV-2 ORF6 protein.

Experimental animal studies evaluating the role of ORF6 protein in SARS-CoV-2 pathogenicity are limited in previous reports. Using K18 human ACE2 transgenic mice, we demonstrated that 50% of the mice infected with ORF6-deficient virus survived while all wild type virus infected mice succumbed to infection (21). The reduced virulence of ORF6-deficient virus infected mice also mirrored in their lower virus loads in nasal turbinate and lungs and reduced lung lesion score compared to wild type virus infected animals (21). Another study in Syrian golden hamsters inoculated with ORF6-deleted virus showed significantly reduced body weight loss and early recovery when compared to animals infected with wild type virus, but similar virus titers were recovered from the nasal turbinates and lungs of both infected groups (13).

Here, we used domestic cat model to study the role of ORF6 in SARS-CoV-2 pathogenesis. This animal model had previously reproduced several characteristics of the virus replication, clinical presentation and respiratory lesions of COVID-19 disease in humans (27, 28, 36). In our current study, we demonstrated that cats inoculated with rWA1 were depressed and had an elevated body temperature; in contrast, rWA1ΔORF6 infected animals were subclinical and remained afebrile. Both inoculated groups, however, lost or maintained steady body weight, whereas the mock-infected cats gained body weight throughout the study. The rWA1ΔORF6 infected cats showed reduced virus replication in nasal turbinate, trachea, and lungs, and lower virus shedding in nasal and oropharyngeal swabs and bronchoalveolar lavage fluid, in comparison to rWA1 infected animals. Histological examination of lungs also revealed a reduced lung pathology in rWA1ΔORF6 infected cats, compared to rWA1 infected animals. Collectively, these findings showed an attenuated phenotype of rWA1ΔORF6 in cats, indicating a role for ORF6 on pathogenesis and the host range of SARS-CoV-2 in felids.

We further investigated the transcriptional response to rWA1 and rWA1ΔORF6 virus infection in the nasal turbinate of cats. The transcriptome of nasal turbinate highlighted the processes underlying the attenuation of rWA1ΔORF6 SARS-CoV-2 virus during infection. We observed marked differences in DEGs in nasal turbinates of cats infected with rWA1 and rWA1ΔORF6 viruses. This points to the fact that the ORF6 gene plays a significant role in viral pathogenesis. Studies have reported that the ORF6 gene disrupts interferon signaling by disrupting the nuclear translocation of IRF3 and STAT1 (15), however, Li and colleagues have reported otherwise that ORF6 is not sufficient to counteract IFN induction during viral infection (39). In this study, we found that type I IFN signaling was significantly upregulated in rWA1ΔORF6 infected cats as compared to rWA1 infected cats, which is consistent with our *in vitro* cell culture experiments, as evidenced by the upregulation of IFN-β and IRF3 promoter activation in luciferase assays. This could be due to significantly up-regulation of the master regulator of ISGs and RSAD2, in rWA1ΔORF6 infected cats compared to rWA1 infected cats. In summary, our study demonstrates the inhibitory role of ORF6 protein on host innate immune responses and deletion of ORF6 attenuates SARS-CoV-2 pathogenesis. These findings will have important implication in coronavirus control and prevention.

## MATERIALS AND METHODS

### Ethics statement

The protocols and procedures for generating recombinant viruses were reviewed and approved by the Cornell University Institutional Biosafety Committee (MUA 16373-2). All animals were handled in accordance with the Animal Welfare Act. The study procedures were reviewed and approved by the Institutional Animal Care and Use Committee at Cornell University (IACUC approval number 2020-0064).

### Cells and viruses

Vero E6 (ATCC^®^ CRL-1586™), and Vero E6/TMPRSS2 (JCRB Cell Bank, JCRB1819) were propagated in Dulbecco’s modified eagle medium (DMEM), supplemented with 10% fetal bovine serum (FBS), L-glutamine (2mM), penicillin (100 U.mL^−1^), streptomycin (100 μg.mL^−1^) and gentamicin (50 μg.mL^−1^). HEK293T cells were maintained in complete growth media consisting of Minimum Essential Media (Corning, 10-010-CV), supplemented with 10% FBS, and penicillin (100 U.mL^−1^) and streptomycin (100 μg.mL^−1^). HEK293T-hACE2 (BEI Resources, NR-52511) cells were grown in Dulbecco’s Modified Eagle’s Medium containing 4 mM L-glutamine, 4500 mg per L glucose, 1 mM sodium pyruvate and 1500 mg per L-sodium bicarbonate, supplemented with 10% fetal bovine serum. The cell cultures were maintained at 37°C with 5% CO_2_.

Sendai virus (SeV) (Cantell strain) was propagated in 11-day-old embryonated chicken eggs and titrated using a hemagglutination assay (HA).

The bacterial artificial chromosome (BAC) harboring the whole genome of the ancestral SARS-CoV-2 WA1 (rWA1) was generated as previously described (25). Deletion of accessory protein ORF6 and generation of recombinant SARS-CoV-2 deficient of ORF6 (rWA1ΔORF6) was described previously (21). For virus rescue, Vero E6 TMPRSS2 cells (3×10^5^ per well of 6-well plate) were transfected with 2 µg/well of the respective SARS-CoV-2 BAC DNA (pBAC-WA1 or pBAC-WA1ΔORF6) using Lipofectamine 3000 (ThermoScientific, USA) and incubated at 37 °C. 24 hours later, the culture medium was replaced with complete growth medium supplemented with 5% FBS and incubated for additional 48 hours at 37 °C. Following the development of cytopathic effects, infected cells and supernatants were harvested and labelled as P0. The recombinant viruses were propagated twice in Vero E6 TMPRSS2 cells. The whole genome sequences of the virus stocks were determined to confirm that no mutations occurred during rescue and amplification in cell culture. The titers of virus stocks were determined by plaque assays and end-point dilutions.

### Generation of feline lung cells stably expressing cat ACE2 receptor

Lentiviral plasmid encoding cat ACE2 (pscALPS-cACE2) was obtained from Addgene (#158082). A c-terminal Myc-tag epitope was added to facilitate detection. Lentiviral particles encoding the cACE2 receptor were produced in HEK293T cells. For this, 7.2 x 10^5^ cells were seeded in each well of a 6-well plate. 24 hours later, cells were transfected with the lentiviral packaging vectors psPAX2 (1 µg, Addgene, 12260), and pMD2.G (1 µg, Addgene, 12259), together with the pscALPS -cACE2 transfer vector (1 µg). Lentiviral particles were harvested from the supernatant of transfected cells at 72 hours post-transfection, cleared by centrifugation, aliquoted and stored at -80 °C until use.

Feline lung cells were seeded (7.2 x 10^5^ cells/well) in a 6-well plate and incubated overnight. For transduction, culture media was removed, 1 mL of the lentivirus was added to each well and adsorbed for 2 hours at 37 °C with the plate rocking back and forth every 30 minutes. 2 mL of complete growth media was added to the cells and incubated for 72 hours at 37 °C. For selecting the transduced cells, the media was replaced with complete growth media containing 5 µg.mL^-1^ of puromycin dihydrochloride (Gibco, A1113803). The puromycin selection was continued until the complete death of non-transduced cells and the expression of cACE2 was validated via immunofluorescence and immunoblots through the detection of the c-terminal Myc tag (Myc-Tag-9B11 Mouse mAb, Cell Signaling Technology, 2276).

### Viral replication kinetics

Viral growth kinetics of rWA1 and rWA1ΔORF6 were performed using three different cell lines: Vero E6 TMPRSS2 (120,000 cells per well), Calu-3 (200,000 cells per well), and feline lung cACE2 (250,000 cells per well). Cells were seeded in 12-well plates for 24 hours until they reached 80-90% confluency. Cells were then inoculated with rWA1 and rWA1ΔORF6 (MOI 0.1) and incubated at 4°C for 1 hour for virus adsorption. After that, the inoculum was replaced with 1 mL of complete growth media and incubated at 37°C. Cells and supernatants were harvested at 4-, 8-, 12-, 24-, 48- and 72-hours pi and stored at -80°C. Time point 0 hour was an aliquot of virus inoculum stored at -80°C as soon as inoculation was completed. Virus titers were determined in Vero E6 TMPRRSS2 cells on each time point using end-point dilutions and the Spearman and Karber’s method and expressed as TCID_50_.mL^−1^.

### Plaque phenotype

The viral plaque phenotype was determined in Vero E6 cells. For this, Vero E6 cells (3×10^5^ cells per well) were seeded in 6-well plates for 24 hours. Cells were inoculated with rWA1 and rWA1ΔORF6 (10 plaque forming units [PFU] per well) and incubated at 37°C for 1 hour. Following that, the inoculum was removed, and 2 mL of media containing 2X complete growth media and 0.5% SeaKem agarose (final conc. 1X media 0.25% agarose) was added to each well. Once the agarose polymerized, the plate was transferred to the incubator at 37°C for 72 hours. The agarose overlay was removed, cells were fixed with 3.7% formaldehyde solution for 30 minutes and stained with 0.5% crystal violet solution for 10 minutes at room temperature. The plaque size was quantified using a Keyence BZ-X810 Microscope.

### Luciferase reporter assays

The ability of the SARS-CoV-2 ORF6 protein to modulate innate immune pathways was investigated using luciferase reporter assays. Initially, a lentiviral plasmid encoding the SARS-CoV-2 ORF6 protein (pLVX-EF1alpha-SARS-CoV-2-orf6-2xStrep-IRES-Puro, Addgene # #141387) was used in the luciferase reporter assays. For this, HEK293T cells were seeded in 24-well plates at 2×10^5^ cells/mL for 24 hours and transfected with pIFN-β-Luc, pNF-κB-Luc, pIRF3-Luc, pPRDII-Luc, or pPRDIII-Luc (200 ng/well) and pRN-Luc (50 ng) reporter plasmids in combination with ORF6 protein expressing plasmid or a pLVX empty vector (250 ng) using Lipofectamine 3000 (Invitrogen, L3000001). At 24 hours post-transfection, cells were stimulated with SeV (Cantell strain, 100 hemagglutination units/well) or TNF-α (25 ng/well) or transfected with 1 µg of poly(I:C) for 12 hours. After stimulation cells were lysed and luciferase activity was measured using the dual luciferase assay (Promega, E2940). Luminescence was measured using a luminometer plate reader (BioTek Synergy LX Multimode Reader). The ratio of luminescence obtained from the target reporters (pIFN-β-Luc, pNF-κB-Luc, pIRF3-luc, pPRDII-luc or pPRDIII-luc) to luminescence from the control Renilla reporter (pRN-Luc) was calculated to normalize the transfection efficiency. Then, the relative pIFN-β-, pNF-κB-, pIRF3-, pPRDII- or pPRDIII-driven luciferase activity was calculated as fold changes over the unstimulated cells. Confirmation of the ORF6 expression in HEK293T transfected cells was performed by Western blot using an antibody against Strep-tag II (GenScript).

Furthermore, the luciferase reporter assay was performed using the two recombinant viruses rWA1 and rWA1ΔORF6. For that, HEK293T-hACE2 cells were seeded in 24-well plates (2×10^5^ cells/mL). After 24 hours, cells were co-transfected with IFN-β or NF-κB Firefly reporter plasmids (200 ng/well) and the control Renilla reporter (pRN-Luc) plasmid (50 ng/well) for 6 hours. Cells were then infected with rWA1 or rWA1ΔORF6 (MOI = 3) for 12 hours. After that, cells were stimulated with SeV (Cantell strain, 100 hemagglutination units/well) or TNF-α (25 ng/well) for 8 hours, cell lysates were harvested, and the luciferase activity was measured using a luminometer. The ratio of luminescence obtained from the target reporters (IFN-β-Luc or NF-κB-Luc) to luminescence from the control Renilla reporter pRN-Luc was calculated to normalize the transfection efficiency. The relative IFN-β- or NF-κB-driven luciferase activity was calculated as fold changes over the unstimulated cells.

### Animals and housing

A total of 18 domestic cats (*Felis catus*) (9 male and 9 female) of 15-18 months of age were obtained from ClinVet (Waverly, NY, USA). The animals were donated to Cornell University to support the reduction of animal use in research. They were housed individually in Horsfall HEPA-filtered cages in the animal biosafety level 3 (ABSL-3) facility at the East Campus Research Facility (ECRF) at Cornell University. Foods and water were provided *ad libitum*.

### Experimental design and sample collection

After acclimation, nasal swabs and blood samples were collected from all animals and tested by rRT-PCR for SARS-CoV-2 and virus neutralization using SARS-CoV-2 B.1 and Omicron BA1.1 lineages, which confirmed that the animals were negative for SARS-CoV-2 (28). On day 0, cats were anesthetized and inoculated intranasally with 1 mL virus suspension (0.5 mL per nostril) containing 5×10^5^ PFU of SARS-CoV-2 rWA1 (n = 6) or rWA1ΔORF6 (n = 6). The control animals (n = 6) received 1 mL of Vero E6 cell culture supernatant intranasally. Body weight and body temperature were recorded daily until day 5 post-inoculation (pi). Oropharyngeal (OPS) and nasal swabs (NS) were collected under sedation (dexmedetomidine) on days 0, 1, 3, and 5 pi using sterile swabs and placed in 1 mL viral transport medium (VTM Corning, Glendale, AZ, USA) and stored at -80°C until used. Three cats from each group were humanely euthanized on days 3 and 5 pi. Necropsy was performed, and bronchoalveolar lavage fluid, tissues such as nasal turbinate, tonsils, retropharyngeal lymph nodes, trachea, lungs, mediastinal lymph node, heart, liver, spleen, kidney, small intestine, and mesenteric lymph node were collected for rRT-PCR and virus isolation. Besides, tissue sections of approximately 0.5 cm from nasal turbinate, trachea, and lungs were collected in 10% neutral buffered formalin (≥20 volumes fixative to 1 volume tissue) for histological examination and *in situ* hybridization (ISH). After 72 hours of fixing, formalin-fixed tissues were transferred to 80% ethanol and processed for histology.

### Nucleic acid isolation and real-time reverse transcriptase PCR (rRT-PCR)

A 10% (w/v) tissue homogenate was prepared using a stomacher (one-speed cycle of 60s, Stomacher® 80 Biomaster). Then, the tissue homogenate supernatant was centrifuged at 2000 x g for 10 min. Viral nucleic acid was extracted from 200 µL of OPS, NS, BALF, and tissue homogenate using the MagMax Core extraction kit (Thermo Fisher, Waltham, MA, USA) and the automated KingFisher Flex nucleic acid extractor (Thermo Fisher, Waltham, MA, USA). The rRT-PCR was performed using the EZ-SARS-CoV-2 Real-Time RT-PCR assay (Tetracore Inc., Rockville, MD, USA) as described earlier (28). An internal inhibition control was included in all reactions. Positive and negative amplification controls were run side-by-side with test samples. A standard curve was performed by using ten-fold serial dilutions from 10^0^ to 10^−8^ of virus suspension containing 10^6^ TCID_50_.mL^-1^ of the SARS-CoV-2 strain. Relative viral genome copy numbers were calculated based on the standard curve and determined using GraphPad Prism 9 (GraphPad, La Jolla, CA, USA). The amount of viral RNA detected in samples was expressed as log (genome copy number) per mL.

### Virus isolation and titrations

The SARS-CoV-2 rRT-PCR positive samples were selected for virus isolation and endpoint titration under Biosafety Level 3 (BSL-3) conditions at the Animal Health Diagnostic Center (ADHC) Research Suite at Cornell University. To this end, Vero E6 TMPRSS2 cells seeded in 96-well plates for 24 were inoculated with serial 10-fold dilutions of the OPS, NS, BALF and tissue homogenates. After 48 hours, cells were fixed with 10% neutral buffered formalin, and IFA was performed as described previously (28). The limit of detection (LOD) for infectious virus titration is 10^1.05^ TCID_50_.mL^−1^. Virus titers were determined at each time point using end-point dilutions and the Spearman and Karber’s method and expressed as TCID_50_.mL^−1^.

### *In situ* RNA detection

Formalin-fixed tissues (nasal turbinate, lungs, and trachea) collected from control and SARS-CoV-2 infected cats at days 3 and 5 pi were embedded in paraffin, sectioned at 5 µM and processed for *in situ* hybridization (ISH) using the RNAscope® ZZ probe technology (Advanced Cell Diagnostics, Newark, CA). The RNAscope^®^ 2.5 HD Reagents–RED kit (Advanced Cell Diagnostics) and a probe targeting SARS-CoV-2 RNA S (V-nCoV2019-S probe ref # 848561) were used in the ISH. A probe targeting feline host protein peptidylprolyl isomerase B (PPIB) was used as a positive control (Advanced Cell Diagnostics cat # 455011). A probe targeting the DapB gene from Bacillus subtilis strain SMY was used as a negative control (Advanced Cell Diagnostics cat # 310043).

### Histology

Formalin-fixed tissues (nasal turbinate, trachea, and lungs) were embedded in paraffin and sectioned at 5µM diameter following standard procedures. The tissue sections were processed for the routine hematoxylin and eosin (H&E) staining and histological examination.

### Bulk RNA-seq library prep and sequencing

To understand the effect of ORF6 gene deletion from wild-type SARS CoV-2 (WA-1) genome on host tissue responses, we performed bulk RNA-seq from nasal turbinate tissue collected at 3 dpi. Total RNA was extracted from nasal turbinate using an RNA extraction kit (MagMax CORE Extraction) and after on-column DNAse treatment, the total RNA integrity score (RIS) with determined with QIAxcel advanced capillary electrophoresis system (Qiagen), according to the instructions. The NGS cDNA library was prepared by using KAPA RNA HyperPrep Kit with RiboErase Kit (HMR) (Roche, USA) and sequenced with Illumina NovaSeq platform with S1-per lane producing 800 million reads100bp Single-Reads (Illumina, USA).

### Bulk RNA-seq data prep-processing

The high-quality, clean data from raw reads were obtained by removing reads with adapters and below Q30 value (FastQC and Trimmomatic). RNA-STAR was used for sequence alignment to reference cat genome (GCF_000181335.3_Felis_catus_9.0_genom.fasta), and the transcripts were quantified with annotated cat genome (cat GCF_000181335.3_Felis_catus_9.0_genome.gtf). The obtained reads per gene and features count were used to obtain the gene expression across different groups. DESeq2 was used to compare differentially expressed genes (DEGs) in different groups (mock, rWA1ΔORF6, and WA-1). Only the genes with log fold change (logFC) ≥2 as biologically significant and statistical (p<0.05) significant were considered as DEGs. The heatmap showing the hierarchical clustering of DEGs across different samples and groups was generated to facilitate the comparison of expression profiles between different groups.

### Gene ontology and pathway enrichment analysis

The DEGs in rWA-1, rWA1ΔORF6, and mock groups were used for gene ontology (GO) biological processes (BP), GO molecular functions (MP), and GO cellular responses (CR) using EnrichR bioinformatic resources (https://maayanlab.cloud/Enrichr/). To identify the fold enrichment of specific genes in cell signaling pathways the statistical overrepresented test (Binomial, P ≤ 0.05) was performed by using GCF_000181335.3_Felis_catus_9.0_genom.fasta as reference. The individual bar graphs for pathway enrichment were created using GraphPad Prism software (version 9.0.1).

### Gene expression verification by real-time quantitative PCR (rRT-PCR)

Total RNA extracted from nasal turbinate of cats at 3 dpi for RNA-seq study was used in gene verification studies. cDNA was synthesized by ProtoScript® II First Strand cDNA Synthesis Kit (NEB). rRT-PCR was performed using PrimeTime Gene Expression Master Mix (IDT) following the manufacturer’s instructions. The following forward and reverse primers and probes were used; IFIH1: forward CACAGGAGTGACTGTCTCATATTC, reverse TTAGACGGCCTCCAAGATTTC, probe TGCCCACACTCAAAGTTCAGGGAT; BST2: forward CCTGCAACAAGACTTTGGTAAC, reverse TGCTTCAACTCCTCAATCTCTC, probe TTCTCCATCTCCAGGGAAGCCAAC; ATF3: forward CTGCAGAAAGAGTCGGAGAAG; reverse CCGATGAAGGTTGAGCATGTA, probe TTCAGTTCGGCATTCACGCTCTCC; TRIM6: forward AAGTGAGTTCTGGACCCTAAAG, reverse GGACATCCGTCAGTTCTCTAAA, probe TGCCCACCAAGCTGAAGAGTATGT; IDO1: forward TGACCTCAAAGACCACAAGTC, reverse TGGCAAGACCTCACAAACA, probe TATGTGTGGAACCAAGGCGGTGAA; SAMHD1: forward TGTGGAATGGACGCATGAA, reverse CTTCAGGGATGAGACCGTAATG, probe AGGGCTCAGTGAAGATGTTTGAGCA; S100A9: forward GCATTGAGACCATCATCAACATC, reverse CACCAGCTGTTTCAGTTCTTTC, probe TGGAGCACCCGGACAAACTCAA; IFIT2: forward CTATGGCAACTTCCAGCTCTAC, reverse GTCTCCTTTGAGTCCTGCTTTAT, probe CGAGGACAGAGCCATCCACCATTT; RSAD2: forward TGGTGAGGTTCTGCAAAGAG, reverse AGATGGCAAGGATGTCCAAATA, probe CGCTGGTTCAGGACTTACGGTGAA; GAPDH: forward AAGGCTGAGAACGGGAAAC, reverse ATACTCAGCACCAGCATCAC, probe TGGAAAGCCCATCACCATCTTCCA. All primers were first tested for their specificity and efficiency using serial 5-fold dilutions of cDNA prepared from pooled nasal turbinate RNA of SARS-CoV-2-inoculated cats.

### Statistical analysis and data plotting

Statistical analysis was performed by 2-way analysis of variance (ANOVA) followed by multiple comparisons. Statistical analysis and data plotting were performed using the GraphPad Prism software (version 9.0.1). Figures 1A and 4A were created with BioRender.com.

## Acknowledgments

We thank the Center for Animal Resources and Education (CARE) staff and Cornell Biosafety team for their support. We thank Dr. Cara Mitchel and all staff at Clinvet for their support and donation of the animals. This work was funded by the National Institute of Health (NIH) and National Institute of Allergy and Infectious Diseases (NIAID) (grant no. R01AI166791-01).

